# Antibiotic treatment induces activation of microglia and impairment of cholinergic gamma oscillations in the hippocampus

**DOI:** 10.1101/2021.07.30.454300

**Authors:** Gürsel Çalışkan, Timothy French, Markus M. Heimesaat, Ildiko Rita Dunay, Oliver Stork

**Affiliations:** Institute of Biology, Otto-von-Guericke University, Magdeburg, Germany; Center for Behavioral Brain Sciences, Magdeburg, Germany; Institute of Inflammation and Neurodegeneration, Medical Faculty, Otto-von-Guericke-University, Magdeburg, Germany; Institute of Microbiology, Infectious Diseases and Immunology, Charité – University Medicine Berlin, Berlin, Germany

**Author notes:** **Corresponding Author** Dr. Gürsel Çalışkan, Institute of Biology, Otto-von-Guericke-University, Leipziger Str. 44, Haus 91, Magdeburg, 39120, Germany. Shared first authorship (Gürsel Çalışkan, Timothy French). Equally contributed last authorship (Ildiko Rita Dunay, Oliver Stork).

**Keywords:** Antibiotics, immune function, microglia, hippocampus, synaptic transmission, gamma oscillations

## Abstract

Antibiotics are widely applied for the treatment of bacterial infections, but their long-term use may lead to gut flora dysbiosis and detrimental effects on brain physiology, behavior as well as cognitive performance. Still, a striking lack of knowledge exists concerning electrophysiological correlates of antibiotic-induced changes in gut microbiota and behavior. Here, we investigated changes in the synaptic transmission and plasticity together with behaviorally-relevant network activities from the hippocampus of antibiotic-treated mice. The prolonged antibiotic treatment led to a strong reduction of myeloid cell pools in bone marrow, circulation and those surveilling the brain. Upon antibiotic treatment, circulating Ly6C^hi^ inflammatory monocytes adopted a proinflammatory phenotype with increased expression of CD40 and MHC II. In the central nervous system, microglia displayed typical signs of activation with elevated CD40 and MHC II expression, as well as increased IL-6 and TNF production. Concomitantly, we detected a substantial reduction in the synaptic transmission in the hippocampal CA1 after antibiotic treatment. In line, carbachol-induced cholinergic gamma oscillation were reduced upon antibiotic treatment while the incidence of hippocampal sharp waves was elevated. These changes were associated with the global changes in the expression of neurotrophin nerve growth factor and inducible nitric oxide synthase, both of which have been shown to influence cholinergic functions. Overall, our study demonstrates that antibiotic-induced changes in the gut microbiome and immune cell function are associated with a drastic reduction in the synaptic transmission and gamma oscillations in the hippocampus, a brain region that is critically involved in mediation of innate and cognitive behavior.

## 1. Introduction

Millions of humans are being prescribed antibiotics (Abx) every day (Suda et al., 2014). However, accumulating evidence from human and rodent studies indicate that the long-term use of antibiotics can lead to detrimental effects on hematopoiesis, brain physiology, innate behavior, and cognitive functions (Desbonnet et al., 2015; Fröhlich et al., 2016a; Josefsdottir et al., 2017; Möhle et al., 2016b). This is due, at least in part, to the decimation of microbiota that colonize our intestinal tract and the disruption of normal gut microbiota-to-brain signaling.

Converging evidence points toward a critical role of microbiota in regulation of the crosstalk between peripheral immune cells and the central nervous system (CNS) (Baruch et al., 2015; Schwartz et al., 2013). We have previously shown that prolonged antibiotic-induced microbial depletion reduces hippocampal neurogenesis and memory retention. Moreover, we demonstrated that these changes are partially mediated by circulating Ly6C^hi^ monocytes as adoptive transfer of these cells from naïve mice could rescue the observed neurogenesis and memory decreases (Möhle et al., 2016b). Monocytes and monocyte-derived macrophages comprise a fundamental leukocyte subset of the innate immune response with multifaceted roles in maintenance of host tissue homeostasis (Biswas et al., 2015; Hammond et al., 2014; Möhle et al., 2016a). Depending on the environment, infiltrating monocytes and CNS resident microglia support diverse functions ranging from inducing to resolving neuroinflammation (Cardona et al., 2006; Michell-Robinson et al., 2015; Shechter and Schwartz, 2013). However, the details of this crosstalk in the homeostatic regulation of normal CNS function remains largely elusive.

One hub region mediating distinct aspects of cognitive function as well as affective behavior is the hippocampus (Çalışkan and Stork, 2018; Maren et al., 2013; Möhle et al., 2016b). Changes in the synaptic transmission and plasticity in hippocampal circuitries have been studied for decades as correlates of these functions (Citri and Malenka, 2008). Indeed, abnormal hippocampal synaptic transmission or plasticity has been reported in numerous neuropsychiatric, neurodegenerative and/or neurodevelopmental conditions with aberrant neurocognitive functions (Annamneedi et al., 2018; Connor et al., 2011; Maggio and Segal, 2011; Rowan et al., 2003). Despite the convincing evidence for the importance of healthy/intact gut microbiota for normal emotional and executive behavior (Bercik and Collins, 2014; Desbonnet et al., 2015; Heijtz et al., 2011; Mayer, 2011; Möhle et al., 2016b; Mostafa and Miller, 2014; Neufeld et al., 2011; Sarkar et al., 2020), to date, there is no direct evidence for the impact of antibiotic-induced gut dysbiosis on hippocampal synaptic transmission and its plasticity.

The hippocampus generates diverse oscillatory rhythms that represent distinct behavioral and cognitive states (Buzsaki, 2004). Specifically, gamma oscillations (30-100 Hz) serve as a common oscillatory mechanism not only for memory encoding/retrieval (Fell and Axmacher, 2011; Montgomery and Buzsaki, 2007), but also for affective behaviors such as fear and anxiety (Headley and Paré, 2013). In the hippocampus, generation and sustainment of gamma oscillations are strongly dependent on cholinergic level both *in vitro* and *in vivo* (Caliskan et al., 2015; Fisahn et al., 1998; Vandecasteele et al., 2014). Of note, neurotrophic factors such as nerve growth factor (NGF) are important regulators of cholinergic activity and potentially can influence hippocampal gamma oscillations (Conner et al., 2009). In addition, microglia activity and glia-associated factors such as tumor necrosis factor (TNF) and inducible nitric oxide synthase (iNOS) can modulate hippocampal synaptic transmission or gamma oscillations (Adaikkan and Tsai, 2020; Beattie et al., 2002; Iaccarino et al., 2016; Martorell et al., 2019; Papageorgiou et al., 2016; Ta et al., 2019). On the other hand, under low cholinergic tonus, hippocampal circuits generate sharp wave-ripples both *in vivo* and *in vitro* (Buzsáki, 2015; Maier et al., 2003). These events may represent on-going plasticity in hippocampal circuitries (Çalışkan and Stork, 2018) and are associated with successful memory consolidation (Girardeau and Zugaro, 2011); pathological alterations in their incidence have been observed in association with abnormal memory formation (Çalışkan et al., 2016; Polepalli et al., 2017). To our knowledge, however, no studies have investigated the impact of antibiotic-induced gut dysbiosis on these behaviourally relevant hippocampal network activities to date.

Given the profound impact of antibiotic treatment on cognitive performance and affective behavior (Desbonnet et al., 2015; Fröhlich et al., 2016b; Möhle et al., 2016b), we hypothesized that antibiotic-induced gut dysbiosis might be associated with aberrant changes in the hippocampal synaptic physiology and associated brain rhythms. Accordingly, we elucidated several factors associated with peripheral and central immunoregulation, cytokine and neurotrophin levels in the CNS, and investigated hippocampal synaptic transmission as well as network oscillations upon antibiotic treatment. Our study indicates that gut dysbiosis is associated with strong alterations in the peripheral and CNS immune function and provides first insights into the potential impact of gut dysbiosis on hippocampal synaptic transmission and behaviorally-relevant hippocampal network activities.

## 2. Methods

### 2.1. Animals

C57B/6JBomTac mice were obtained from M&B Taconic, Germany, and bred in the animal facility at the Otto-von-Guericke University Magdeburg (12h light/dark cycle with lights switched on at 19:00 hr with a 30 min dawn phase; food and water ad libitum). All experiments were conducted in accordance with the European and German regulations for animal experiments and were approved by the Landesverwaltungsamt Saxony-Anhalt (Permission Nr. 203.h-42502-2-887 OvGMD; 203.h-42502-2-1206 UniMDOVGU).

### 2.2. Antibiotic Treatment and microbiota assessment

Male mice were group-housed (3 to 6 mice per cage) and treated with broad-spectrum antibiotics (Abx) according to the protocol previously published (Möhle et al., 2016b). The Abx compounds consisting of ampicillin plus sulbactam (1.5□g/L; Pfizer), vancomycin (500 mg/l; Cell Pharm), ciprofloxacin (200 mg/l; Bayer Vital), imipenem plus cilastatin (250 mg/l; MSD) and metronidazole (1 g/l; Fresenius) were dissolved in 1 L autoclaved water. Treatment began with an initial per oral challenge of 200 µL Abx mixture using an oral gavage. Then Abx mixture was applied via the drinking water *ad libitum* and continued until the end of the experiment. Drinking water was switched with fresh Abx twice a week. Mice were relocated to new cages every other day to prevent recolonization from feces.

Fecal samples were taken to monitor antibiosis in antibiotic-treated mice. DNA from fecal samples was extracted as described previously (Heimesaat et al., 2010). Briefly, DNA extracts and plasmids were quantified using QuantiT PicoGreen reagent (Invitrogen) and adjusted to 1 ng/μl. Main bacterial groups abundant in the murine conventional intestinal microbiota were assessed by quantitative RT-PCR with group-specific 16S rRNA gene primers (Tib MolBiol) (Rausch et al., 2013). The number of 16S rRNA gene copies per ng DNA of each sample was determined and frequencies of respective bacterial groups calculated proportionally to the eubacteria amplicon.

### 2.3. Immunology methods

#### 2.3.1 Cell Isolation

Blood immune cells from mice were collected from the vena cava and prepared as previously described (Biswas et al., 2015). Mice were deeply anaesthetized by isoflurane inhalation (CP Pharma) and intracardially perfused with 60 mL sterile phosphate-buffered saline (PBS) prior to organ extraction. To isolate bone marrow cells, the femur and tibia were isolated and surrounding tissue was removed. The bone ends were removed, the bone marrow cells were washed out with FACS buffer using a syringe and a 26□G needle and sieved through a 40 μm strainer. Brains were homogenized in a buffer containing HBSS (Gibco), 1M HEPES (pH 7.3, Thermo Fisher) and 45% glucose before sieving through a 70 µm cell strainer. The homogenate was fractioned on a discontinuous 30-70 % Percoll gradient (GE Healthcare). Immune cells were collected from the 30/70 % Percoll interphase, washed in PBS and stained for flow cytometric analysis.

#### 2.3.2 Flow cytometry

Flow cytometry stainings were performed as previously described (Düsedau et al., 2019). Single Cell suspensions were incubated with an anti-FcγIII/II receptor antibody (clone 93, eBioscience) to block unspecific binding and Zombie NIR™ (Biolegend), a fixable viability dye. Thereafter, cells were stained with fluorochrome-conjugated antibodies against cell surface markers: CD45 (30-F11), CD11b (M1/70), Ly6C (HK1.4), CD40 (HM-40-3), F4/80 (BM8), Ly6G (1A8) and MHCII (M5/114.15.2) all purchased from BioLegend; CD40 (HM-40-3) purchased from eBioscience in FACS buffer at 4□°C for 30□min. Matched FMO controls were used to assess the level of background fluorescence in the respective detection channel.

To measure cytokine production, intracellular staining was performed as previously described (Düsedau et al., 2019). Cells were incubated with an anti-FcγIII/II receptor antibody and Zombie NIR™ (Biolegend). Then surface staining was performed on the cells with antibodies against CD45 (30-F11), CD11b (M1/70), Ly6C (HK1.4) and Ly6G (1A8) in FACS buffer. Cells were then fixed in 4% paraformaldehyde and permeabilized using Permeabilization buffer (Biolegend). Intracellular proteins were stained with antibodies against TNF (MP6-XT22) and IL-6 (MP5-20F3) purchased from eBioscience. Matched isotype controls were used to assess the level of non-specific binding. Flow cytometric analysis was performed on an Attune NxT Flow Cytometer (Thermo Fisher) and analyzed with FlowJo (version 10, FlowJo LLC).

#### 2.3.3 RNA Isolation

To isolate total RNA, samples were homogenized in BashingBeads tubes (Zymo Research) and RNA was isolated using peqGOLD total RNA kit (Peqlab) according to the manufacturer’s instructions.

#### 2.3.4 RT-qPCR

Gene expression was determined using the TaqMan® RNA-to-CT™ 1-Step Kit (Life Technologies), as previously described (French et al., 2019; Lang et al., 2018). TaqMan® Gene Expression Assays (Life Technologies) were used for mRNA amplification of *Tnf* (Mm00443258_m1), *Il6* (Mm00446190_m1), *Ifn*γ (Mm00801778_m1), *Gfap* (Mm01253033_m1), *Cx3cr1* (Mm02620111_s1), *Slc17a7* (Mm00812886_m1), *Slc1a2* (Mm01275814_m1), *Gabra1* (Mm00439046_m1), *Gria1* (Mm00433753_m1), *Gria2* (Mm00442822_m1), *Bdnf* (Mm04230607_s1), *Ngf* (Mm00443039_m1), *Ntf3* (Mm01182924_m1), *Nox4* (Mm00479246_m1), *Cox2* (Mm00478374_m1) and *Nos2* (Mm00440485_m1). Expression of *Hprt* (Mm01545399_m1) was chosen as reference, and target/reference ratios were calculated with the LightCycler® 96 software version 1.1 (Roche). All results were further normalized to the mean of the (naive) group.

### 2.4. Slice Electrophysiology

Slice electrophysiology was performed as described previously (Caliskan et al., 2015; Çalışkan et al., 2019). After 6 weeks of chronic antibiotic treatment, decapitation of mice and brain extraction were performed under deep isoflurane anesthesia. ∼400 µm thick brain slices that contain ventral-to-mid hippocampus were cut in horizontal plane using an angled platform (12° in the fronto-occipital direction) in ice-cold, carbogenated (5% CO_2_ / 95% O_2_) artificial cerebrospinal fluid (aCSF) containing (in mM) 129 NaCl, 21 NaHCO_3_, 3 KCl, 1.6 CaCl_2_, 1.8 MgCl, 1.25 NaH_2_PO_4_ and 10 glucose (pH 7.4, ∼300 mosmol / kg) with a vibrating microtome (Campden Instruments; Model 752) and quickly transferred to an interface chamber perfused with aCSF at 32 ± 0.5 °C (flow rate: 2.0 ± 0.2 mL / min). A minimum of one hour of slice recovery was allowed before starting recordings. Field potentials (FP) were recorded using borosilicate glass electrodes filled with aCSF with a resistance of ∼1 MΩ. FP responses were evoked with a constant current stimulator and a bipolar tungsten wire stimulation electrode (exposed tips: ∼20 µm; tip separations of ∼75 µm; electrode resistance in aCSF: ∼0.1 MOhm). FP signals were pre-amplified using a custom-made amplifier and low-pass filtered at 3 kHz. Signals were sampled at a frequency of 10 kHz and stored on a computer hard disc for off-line analysis (Cambridge Electronic Design, Cambridge, UK).

#### 2.4.1. Evoked field potential recordings

To obtain field excitatory postsynaptic potential (fEPSP) responses from CA (Cornu Ammonis) 3-to-CA1 synapse, the recording electrode was placed at the apical dendrites of area CA1 (Stratum Radiatum: SR) and the bipolar stimulation electrode was placed on the Schaffer collaterals (SC) at the proximal CA1 close to the CA2 subregion. After placing the electrodes, responses were recorded for ten-to-twenty min until they were stabilized (inter-stimulus interval of 30 sec and stimulation duration of 100 µs). Then, an input-output (I-O) curve was recorded using intensities ranging from 10 to 50 µA. This was followed by paired-pulse (PP) recording protocol with intervals ranging from 10 to 500 ms. After PP protocol, baseline responses were recorded for another 20 min and LTP induction was commenced with a train of 100 pulses (100 Hz) repeated 2 times with 20 s interval. This was followed by test pulses recorded for 40 min (0.033 Hz). MATLAB-based analysis tools were used for the analysis of fEPSPs (MathWorks, Natick, MA). For calculation of fEPSP slopes, the slope (V/s) between the 20 and 80% of the fEPSP amplitudes were measured. We also calculated an average baseline transmission rate per slice by dividing each fEPSP slope value with the corresponding FV value followed by averaging these values leading to one transmission rate (ms^-1^) value per slice. Paired-pulse responses were analyzed by dividing the slope of the second fEPSP to the first one. For the analysis of LTP, the data were normalized to baseline responses obtained for 20 min before LTP induction.

#### 2.4.2. Cholinergic gamma oscillations

To induce gamma oscillations, the temperature of interface chamber was set to 35°C and freshly-diluted carbachol (CBh, 5 µM) was applied via continuous bath perfusion. Fifty-to-seventy min after CBh perfusion, three-to-five min recordings were obtained from pyramidal layer of CA3 subregion (stratum pyramidale: SP). Custom-made spike2 scripts were used to analyze gamma oscillations (Cambridge Electronic Design, Cambridge, UK). From each recording 2 min artifact free data was extracted, and power spectra were generated using Fast Fourier Transformation with a frequency resolution of 0.8192 Hz. Peak frequency (Hz) and Integrated power (20-80 Hz; µV^2^) were calculated from the power spectra. Gamma recordings with peak powers lower than 40 µV^2^ and peak frequencies lower than 20 Hz were discarded. For the autocorrelation analysis auto-correlograms were calculated from the 2 min data. The value of the 2nd positive peak of auto-correlogram was used to report the gamma correlation strength of local CA3 gamma oscillations.

#### 2.4.3. Sharp Wave-Ripples

Glass electrodes were placed at the SP of CA1 subregion. Data were recorded for three-to-five min and two min artifact free data were extracted as MATLAB files to be further analyzed using a custom written MATLAB-scripts (MathWorks, Natick, MA). For sharp wave (SW) detection, the data was low-pass filtered at 45 Hz (FFT filter). The SW events were detected with threshold set to 2.5 times the standard deviation (SD) of the lowpass-filtered signal. The minimum interval between two subsequent SW was set to 80 ms. Data stretches of 125 ms centered to the maximum of the sharp wave event were stored for further analysis. The start and the end point of SW was determined by the points crossing the mean of the data. The area under curve (AUC) was calculated using the low pass filtered data using these two points as the start and end of the SW event. Ripples were isolated using band-pass filter at 120-300 Hz (FFT filter). 15 ms before and 10 ms after the maximum of SW event (25 ms) were stored for further analysis of the ripples. The ripple events were detected with threshold set to 3 times the SD of the bandpass-filtered signal. Ripple amplitude was analyzed using triple-point-minimax-determination. Only ripples with lower than 75% difference between falling and rising component were included in the analysis. The time between the through of subsequent ripples were used for calculation of ripple frequencies.

### 2.5. Statistical analysis

Electrophysiological data were statistically analyzed using SigmaPlot for Windows Version 11.0 (Systat software). Before statistical comparison of different groups, normality test (Shapiro-Wilk Test) and equal variance test were performed. I-O and PP curves were statistically compared using two-way repeated measures ANOVA. For comparison of baseline transmission rate per slice Student’s two-tailed t-test was used. For LTP data, normalized data obtained 30-40 min after LTP induction was used for statistical comparison with Mann-Whitney U test. For hippocampal network oscillations, statistical differences were determined by Student’s two-tailed t-test or Mann-Whitney U test. For the statistical comparison of gamma power, log transformed data was used. Data from RT-qPCR, flow cytometry cell populations and cell activation were analyzed by Mann-Whitney U test. All Mann-Whitney U tests were two tailed. Probability values of p < 0.05 were considered as statistically significant. Sample sizes are provided in figure captions (N: Number of mice; n: Number of slices).

## 3. Results

### 3.1. Microbiota depletion upon antibiotic treatment

Adult mice were treated with a broad-spectrum antibiotic cocktail for 4 weeks. This antibiotic cocktail has been shown to effectively deplete the intestinal microbiota (Möhle et al., 2016b). To confirm the eradication of the commensal gut bacterial species, the feces from mice with and without antibiotic treatment (+Abx vs. naïve, respectively) were collected and analyzed by 16S rRNA sequencing. As expected, broad-spectrum antibiotic treatment led to an elimination of the most abundant gut bacterial groups, genera and species (Fig. 1). Of note, all fecal samples derived from gut microbiota of Abx-treated mice were culture-negative for aerobic, microaerophilic and obligate anaerobic bacteria applying both, solid and enrichment media (data not shown).

**Fig. 1.**
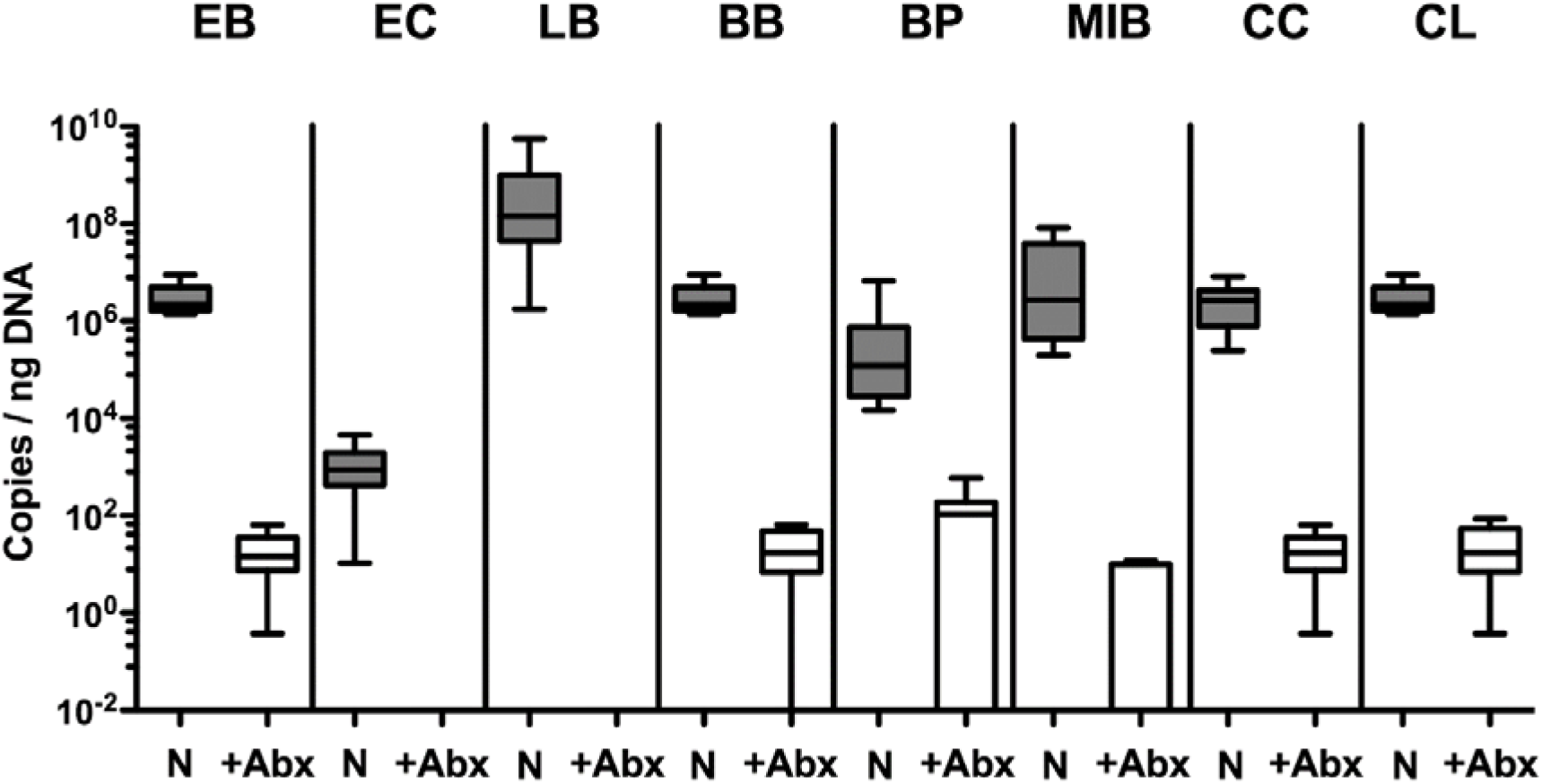
Microbiota depletion upon antibiotic treatment. The gut microbiota composition was quantitatively assessed in fecal samples derived from mice that had been subjected to broad-spectrum antibiotic treatment (+Abx; white boxes, n=10) and from untreated naïve counterparts (N; grey boxes n=10) applying quantitative 16S rRNA-based bacterial real-time PCR amplifying variable regions of the following bacterial groups (expressed as copies per ng DNA): EB, enterobacteria; EC, enterococci; LB, lactobacilli; BB, bifidobacteria; BP, Bacteroides/Prevotella species; MIB, Mouse Intestinal Bacteroides; CC, Clostridium coccoides group; CL, Clostridium leptum group. Box plots represent the 75^th^ and 25^th^ percentiles of the medians (black bar within box).

### 3.2. Reduced frequency of circulating Ly6C^hi^ monocytes and increased immune activation upon antibiotic treatment

Circulating Ly6C^hi^ monocytes exhibit a multifaceted set of functions for the host that have been shown to be beneficial, such as transforming into and replenishing tissue resident DCs or macrophages, or detrimental by exacerbating immunopathology (Cryan and Dinan, 2015; Möhle et al., 2016b; Shi and Pamer, 2011). Thus, we aimed to examine the effect of broad-spectrum Abx treatment on the pool of Ly6C^hi^ monocytes in bone marrow, blood and brain by flow cytometry. Monocytes were distinguished from neutrophils via the surface markers CD11b, Ly6C and Ly6G and from microglia by expression of CD45. Monocytes were gated as CD45^hi^CD11b^+^Ly6C^+^Ly6G^-^ (Fig. 2A, 2C, 2G). Gating of immune cell subsets in the brain that distinguishes recruited myeloid cells from microglia is shown in Fig. 3C. We observed that Abx treatment led to a significant reduction in the frequency of Ly6C^hi^ cells in bone marrow (Fig. 2B; Mann-Whitney U test; p=0.0079), circulating in the blood (Fig. 2D; Mann-Whitney U test; p=0.008) and those infiltrating the brain (Fig. 2H; Mann-Whitney U test; p=0.0079). To investigate if Abx treatment was altering the activation status of the circulating monocytes, we analyzed their expression of MHC II and CD40, markers associated with classically activated M1 immune cells and upregulated upon TLR engagement (Andrade et al., 2005; Sica and Mantovani, 2012). We detected a significant increase in the expression of both MHC II and CD40 in blood Ly6C^hi^ monocytes (Fig 2E-F; Mann-Whitney U test; MHC II, p=0.016; CD40, p=0.016) and increase of CD40 on those infiltrating the brain (Fig 2I-J; Mann-Whitney U test; MHC II, p=0.52; CD40, p=0.021) upon Abx treatment. These results highlight that prolonged Abx treatment associates with a global reduction of the available Ly6C^hi^ monocyte pool and shift towards an activated phenotype. However, MHC II expression remains unaltered when the CNS is infiltrated, suggesting there might be a certain amount of translocation of the remaining gut microbes which promote inflammation locally. We observed an enlarged and inflamed cecum in Abx-treated animals which is comparable to other Abx treatment studies in mice (Ge et al., 2017; Sun et al., 2021). Since the spleen is a primary filter of blood-borne pathogens and antigens and serves as detector of systemic inflammation (Altamura et al., 2001; Cobb et al., 2002), we analyzed the gene expression of TNF and IL-6. Gene expression analysis revealed that Abx treatment led to an increase of both TNF and IL-6 expression compared to naïve animals (Fig 2K; Mann-Whitney U test; TNF, p=0.002; IL-6, p=0.028). These data indicate that microbiota are necessary to maintain the steady-state myeloid populations.

**Fig. 2.**
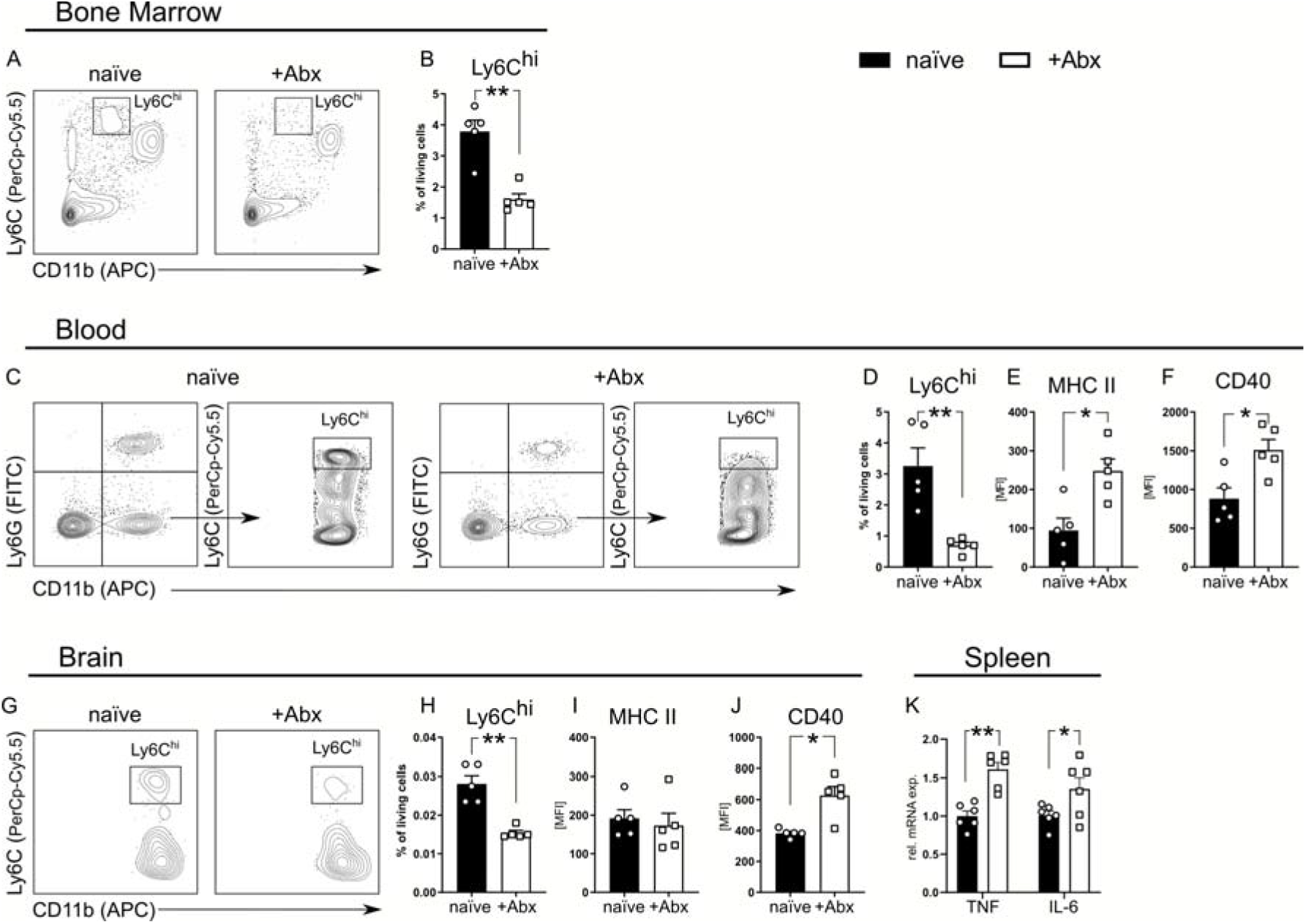
Reduced frequency of Ly6C^hi^ monocytes and increased immune activation upon antibiotic treatment. Immune cells from bone marrow, blood and brain were isolated from naïve (N=5 mice) and Abx-treated (+Abx; N=5 mice) mice and analyzed by flow cytometry. (A, C, G) Representative Ly6C^hi^ monocyte gating strategies for bone marrow, blood and brain. Events were gated on singlets and cells were excluded based on forward and side scatter (FSC/SSC) and viability staining (dead cells).. Ly6C^hi^ monocytes were defined as CD11b^+^Ly6G^-^Ly6C^hi^ (bone marrow, blood) or CD45+CD11b+Ly6G-Ly6Chi (brain, to exclude CD45lowCD11b+ microglia). (B, D, H) The frequency of Ly6C^hi^ monocytes is presented as the percentage of living single cells. (E, F, I, J) The surface expression of MHC II and CD40 was determined on Ly6Chi monocytes in naïve and Abx-treated mice by flow cytometric analysis and presented as median fluorescence intensity (MFI). Spleens were collected from naïve (N=6) and Abx-treated (N=6) mice and were homogenized. (K) Relative mRNA levels were normalized to the mean expression of the naïve control group. Symbols represent individual animals. Data is representative of three independent experiments presented as mean + SEM.

**Fig. 3.**
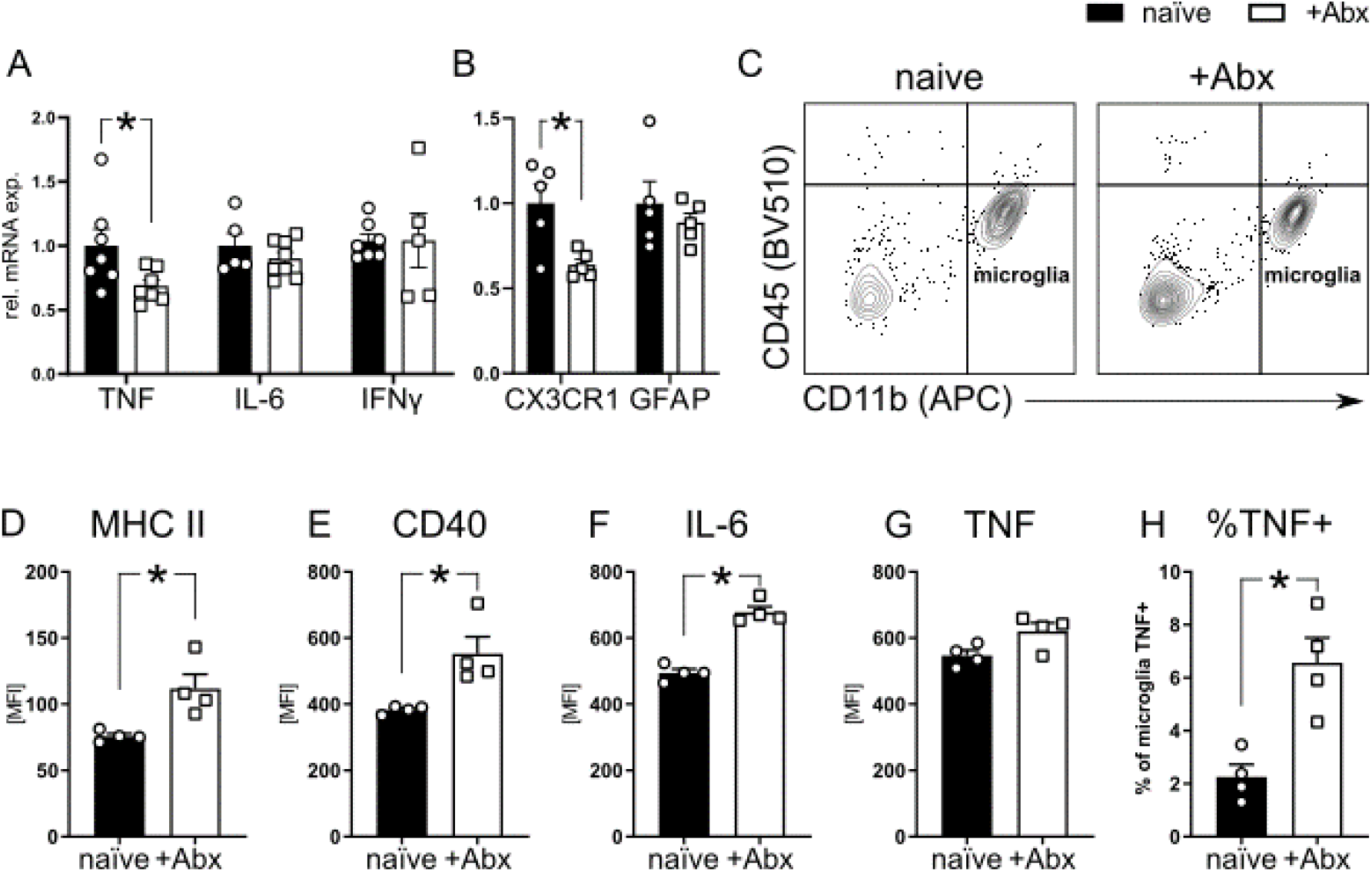
Increased activation of microglia upon antibiotic treatment. Relative expression of mRNA levels of (A) proinflammatory and (B) glial genes in naïve (N=5-7) and Abx-treated (N=5-8). Relative mRNA levels were normalized to the mean of the naïve control group. (C-H) Immune cells from brains of naïve (N=4) and Abx-treated (N=4) were isolated and analyzed by flow cytometry. (C) Representative gating strategy of brain immune cells. Events were gated on singlets and cells were excluded based on forward and side scatter (FSC/SSC) and viability staining (dead cells). CD11b^+^CD45^int^ cells were defined as microglia. The surface expression of (D) MHC II, (E) CD40 and intracellular production of (F) IL-6 and (G) TNF by microglia were quantified and presented as MFI of their respective fluorochrome. (H) The percentage of microglia that are positively producing TNF. RNA was isolated from whole brain homogenate of naïve and Abx-treated (+Abx) mice for RT-qPCR analysis. Symbols represent individual animals. Data is representative of three independent experiments and presented as mean + SEM.

### 3.3. Antibiotic treatment reduces CX3CR1 expression and increases activation of resident microglia

Microglia and recruited myeloid cells are vital for sustaining tissue homeostasis in the CNS (Shechter and Schwartz, 2013). Given the observed reduction of circulating Ly6C^hi^ monocytes and increased splenic TNF and IL-6 expression, we decided to first assess if Abx treatment was also altering the gene expression of proinflammatory markers TNF, IL-6 and IFNγ in whole brain homogenates of naïve and Abx-treated mice. In contrast to the periphery, we detected a decrease of TNF gene expression and no difference in IL-6 or IFNγ gene expression (Fig. 3A; Mann-Whitney U test; TNF, p=0.02; IL-6, p=0.35; IFNγ, p=0.99) in Abx-treated mice. TNF is primarily synthesized by glial cells in the brain (McCoy and Tansey, 2008). To determine if a particular resident glia cell type is altered upon Abx treatment, we measured the gene expression of glial activation markers GFAP (astrocytes) and CX3CR1 (microglia) (Li et al., 2020; Wolf et al., 2013). Following antibiotic treatment, GFAP expression was unchanged whereas expression of CX3CR1 was reduced (Fig. 3B, Mann-Whitney U test; CX3CR1, p=0.03; GFAP, p=0.8), indicating that microglia in particular, are influenced by Abx treatment.

CX3CL1 is mainly secreted by neurons and induces neuroprotective effects via interaction with its receptor CX3CR1 on microglia (Nash et al., 2015). CX3CL1 thereby downregulates the production of proinflammatory mediators such as TNF, NO and superoxides. Based on the observed reduction in CX3CR1, we next examined the activation status of resident microglia (CD11b^+^CD45^int^ cells) via flow cytometry, using MHC II and CD40 as phenotypic markers (Fig. 3C). Upon Abx treatment, expression of MHC II (Fig. 3D; Mann-Whitney U test; p=0.028) and CD40 (Fig. 3E; Mann-Whitney U test; p=0.020) significantly increased. To further investigate the level of microglia activation, cytokine production was assessed. Here, microglia of Abx-treated mice showed a significantly higher IL-6 expression (Fig. 3F; Mann-Whitney U test; p=0.026) compared to naïve mice whereas TNF production remained unchanged (Fig 3G; Mann-Whitney U test; p=0.11). However, there were significantly more microglia producing TNF in Abx-treated mice compared to naïve (Fig. 3H; Mann-Whitney U test; p=0.028). These data imply that the gut microbiome plays an important role in modulating and maintaining microglial function.

### 3.4. Enhanced iNOS expression and reduced NGF upon antibiotic treatment

Microglia are known to upregulate reactive oxygen species (ROS) and reactive nitrogen species (RNS) when activated (Mander and Brown, 2005; Ta et al., 2019). In excessive quantities, ROS can lead to axonal and neuronal loss, for example, in neurodegenerative diseases (Barbeito et al., 2004; Okuno et al., 2005). To investigate if Abx treatment leads to alterations in the gene expression of RNS or ROS producing enzymes, we measured the gene expression of NADPH-Oxidase (*Nox4*), cyclooxygenase 2 (*Cox2*) and inducible nitric oxide synthase (iNOS; *Nos2*), two superoxide and NO producers affiliated with microglial function (Dugan et al., 2009; Ta et al., 2019) via RT-qPCR analysis. Upon Abx treatment, no change was detected in *Nox4* or *Cox2* gene expression whereas *Nos2* expression was significantly increased (Fig. 4A; Mann-Whitney U test; *Nox4*, p=0.99; Cox2, p=0.27; iNOS, p=0.0079). Upregulation of iNOS along with microglia activation has been linked to disturbances in fast neuronal network oscillations underlying perception, attention and memory (Ta et al., 2019). Moreover, we previously demonstrated (Möhle et al., 2016b) that the same antibiotic cocktail disrupts learning and memory. Thus, we hypothesized that Abx treatment may lead to dysregulation of crucial neurotrophins, and proteins involved in synaptic homeostasis. To determine if Abx treatment led to changes in expression of neurotrophins, we measured the mRNA levels of brain-derived neurotrophic factor (BDNF), nerve growth factor (NGF) and neurotrophin-3 (NTF3). There were no gene expression changes observed in BDNF or NTF3 between the groups whereas NGF gene expression was diminished in Abx-treated brains (Fig. 4B; Mann-Whitney U test; *Bdnf*, p=0.41; *Ngf*, p=0.042, *Ntf3*, p=0.52). To explore whether microglial activation and increased *Nos2* expression result in changes to excitatory or inhibitory signal transduction, expression levels of receptors and transporters involved in glutamatergic (Glur1/2, VGLUT1, EAAT2) and GABAergic signaling (GABA_A_α1) were determined. When looking at the gene expression for all synaptic markers, we observed no change in expression after Abx treatment (Fig. 4C; Mann-Whitney U test; p>0.8 for all).

**Fig. 4.**
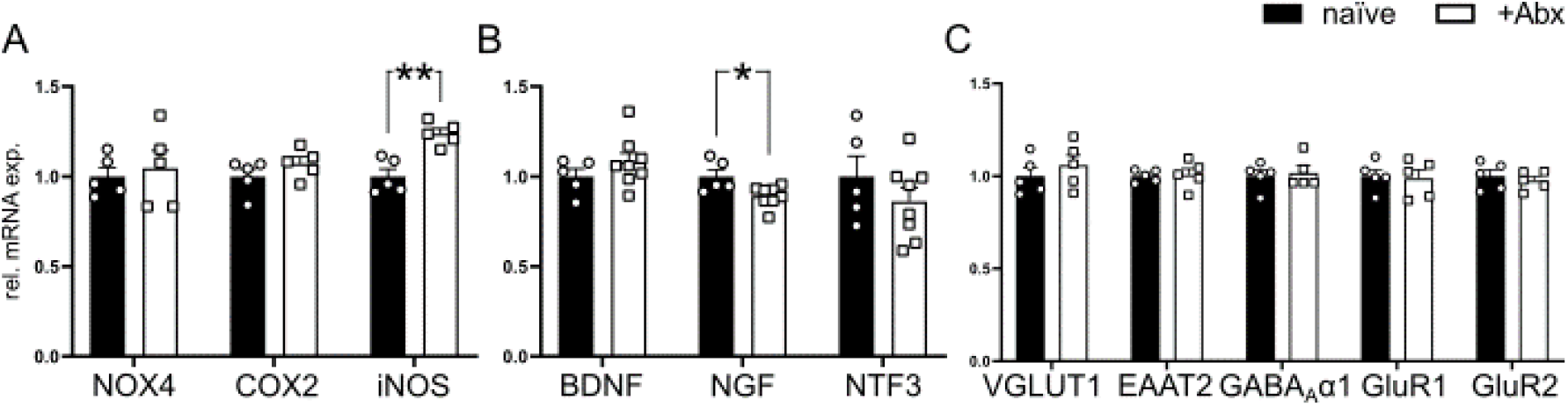
Increased iNOS and decreased NGF gene expression upon antibiotic treatment. RNA was isolated from whole brain homogenate of naïve and Abx-treated (+Abx) mice for RT-qPCR analysis. Relative expression of mRNA levels of (A) reactive species, (B) neurotrophins and (C) synaptic genes in naïve (N=5) and +Abx (5-8) mice. Relative mRNA levels were normalized to the mean of the naïve control group. Symbols represent individual animals. Data is representative of three independent experiments. Data is presented as mean + SEM.

### 3.5. Reduced hippocampal synaptic transmission upon antibiotic treatment

Due to the well-established role of cytokines and neurotrophins in sustaining synaptic transmission and plasticity (Albensi and Mattson, 2000; Beattie et al., 2002; Conner et al., 2009; Curran and O’Connor, 2003; Hoshino et al., 2017; Leal et al., 2017), we assessed the changes in synaptic physiology at the hippocampal CA3-to-CA1 synapse upon chronic Abx treatment. We observed a profound decrease in the baseline transmission evident by lower dendritic fEPSP responses to increasing stimulation strengths (Fig. 5A). We detected a strong stimulation intensity x fEPSP slope interaction (Two-way repeated measures ANOVA; F(6, 27) = 4.569, p < 0.001) with significant reduction at 20 µA to 50 µA stimulus intensities (Fisher’s LSD post hoc comparison; 20 µA: p = 0.25; 30 µA: p = 0.012; 40 µA: p = 0.008; 50 µA: p = 0.008). On the other hand, we observed no Abx treatment effect (Fig. 5B; Two-way repeated measures ANOVA; F(1, 21) = 4.569, p < 0.001) for the presynaptic fiber volley (FV) amplitudes suggesting a potential Abx-induced alteration restricted to the postsynaptic part. Comparison of average baseline transmission rates per slice (see methods), also showed a reduced baseline transmission rate (Fig. 5C; Student’s two-tailed t-test; T(21) = 2,156, p = 0.043) suggesting a reduced efficacy of synaptic transmission at this hippocampal synapse. Assessment of short-term plasticity via paired-pulse protocol revealed no statistical differences after Abx treatment (Fig. 5D; Two-way repeated measures ANOVA; F(1, 27) = 1.378, p < 0.251). Last, we assessed long-term potentiation with a HFS protocol (see methods). Both control and Abx-treated mice showed a strong potentiation which stayed stable up to 40 min after HFS (Fig. 5E). Comparison of normalized fEPSP slope values 30-40 min after HFS showed no effect of Abx treatment (Fig. 5F; Mann-Whitney U test; p = 0.075). These data suggest that chronic Abx treatment results in a profound decrease in the baseline synaptic transmission at the hippocampal CA3-CA1 synapse without altering synaptic plasticity.

**Fig 5.**
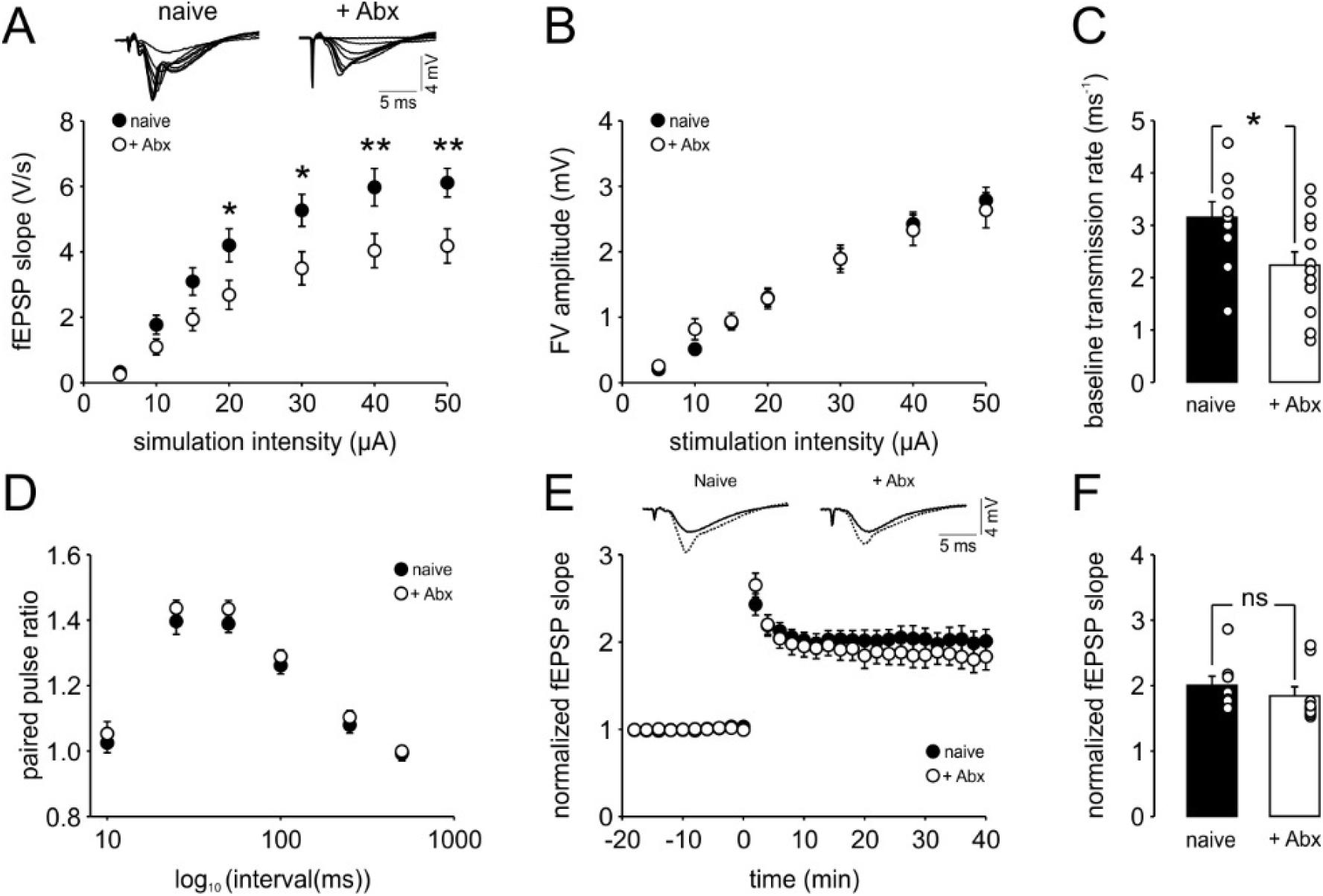
Reduced baseline transmission at the Schaffer collateral (SC)-CA1 synapse after chronic antibiotic treatment (+Abx). **(A)** Input-output curve showing reduced CA1 field excitatory postsynaptic potentials (fEPSP) in +Abx mice (N=5 mice, n=13 slices) in comparison to naive mice (N=3 mice, n=10 slices). **(B)** Input-output curve showing unaltered presynaptic fiber volley amplitudes in +Abx mice (N=5 mice, n=19 slices) in comparison to naive mice (N=3 mice, n=10 slices). **(C)** Summary graph illustrating a reduced baseline transmission rate (averaged fEPSP slope / FV amplitude values per slice) at the SC-CA1 synapse after Abx treatment (Naive mice: N=3 mice, n=10 slices; +Abx mice: N=5 mice, n=13 slices). **(D)** Short-term plasticity is not altered in +Abx mice evident by the unchanged paired pulse ratios (Naive mice: N=3 mice, n=10 slices; +Abx mice N=5 mice, n=19 slices). **(E)** Increase in the fEPSP responses upon high frequency stimulation (HFS) of the SC in the +Abx mice are comparable to the control mice indicating no change in long-term plasticity (LTP). **(F)** Analysis of normalized fEPSP values obtained during 30-40 min after HFS reveal no significant alteration between the naive (N=4 mice, n=8 slices) and +Abx (N=5 mice, n=9 slices) mice. Data are presented as mean ± SEM.

### 3.6. Reduced hippocampal gamma oscillations upon antibiotic treatment

In order to assess the impact of the reduced hippocampal synaptic transmission on behaviorally-relevant activities in the hippocampus, first, we recorded cholinergic gamma oscillations in the CA3 subregion. Gamma oscillations appeared after ten-to-fifteen min and stabilized after 50 min after CBh perfusion (Fig. 6A). Analysis of the power spectra revealed a shift in the peak frequency of gamma oscillations (Fig. 6B-C; Mann-Whitney U test; p < 0.001) and strong reduction in the gamma power (Fig. 6C-D; Student’s two-tailed t-test; T(28) = 2.591, p = 0.015). Next, we measured local synchronization of gamma oscillations and found no significant change in the gamma correlation upon Abx treatment (Fig. 6E-F; Mann-Whitney U test; p = 0.271). Overall, gamma oscillations appear to be reduced in power upon chronic Abx treatment.

**Fig 6.**
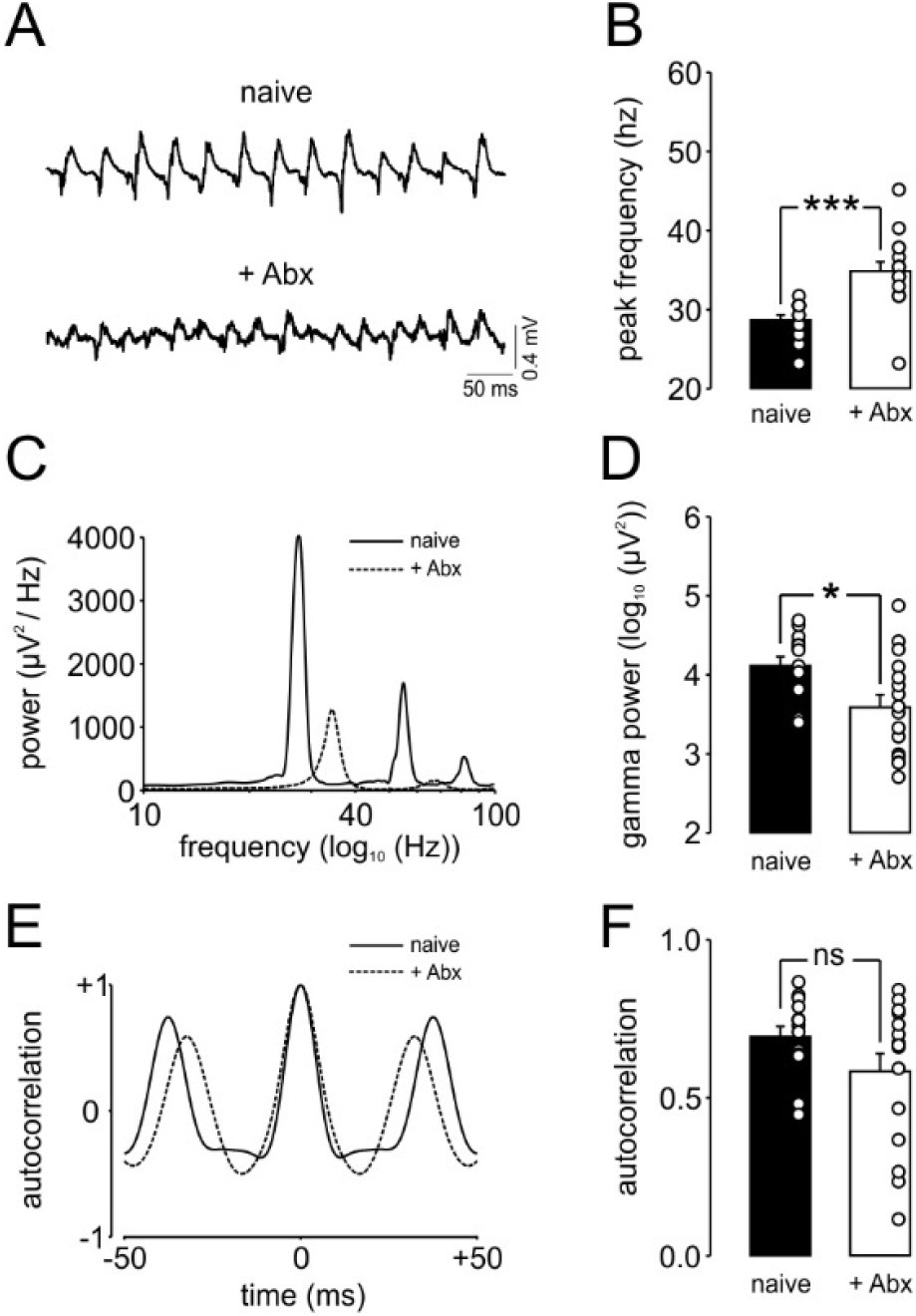
Reduced hippocampal cholinergic gamma oscillations after chronic antibiotic treatment. **(A)** Representative carbachol (CBh, 5 µM) induced local field potential (LFP) traces from hippocampal CA3 of +Abx (N=5 mice, n=16 slices for all parameters) and naive (N=4 mice, n=14 slices for all parameters) mice. **(B)** Summary graph illustrating an increased gamma peak frequency in +Abx mice. **(C)** Representative power spectra illustrating a shift in the main gamma peak frequency and a reduction in gamma-range oscillation power in the hippocampal CA3 of +Abx mice. **(D)** Summary graph illustrating an increased gamma power (20-80 Hz) in the hippocampal CA3 of +Abx mice. Representative auto-correlograms of CA3 LFP gamma oscillations. **(E)** Representative auto-correlograms illustrating a shift in the 2^nd^ positive peak indicating a reduced duration of gamma cycles, thus, increased gamma peak frequency, in +Abx mice **(F)** Summary graph showing no significant change in the amplitude of the 2^nd^ peak of the gamma auto-correlograms of +Abx mice in comparison to naive mice. Data are presented as mean ± SEM.

### 3.7. Increased incidence of sharp waves upon antibiotic treatment

Horizontal or transverse-like slices obtained from the ventral-to-mid portion of the hippocampus generates spontaneous sharp wave-ripple (SW-R) activity (Caliskan et al., 2015; Çalışkan et al., 2016; Maier et al., 2003). Both *in vivo* and *in vitro*, these events have been associated with emotional and spatial memory consolidation (Çalışkan et al., 2016; Girardeau et al., 2017, 2009). Thus, we assessed the effect of chronic Abx treatment on spontaneous SW-R activity in the CA1 subregion (Fig. 7A-B). We found a specific increase in the incidence of SW (Fig. 7C; Student’s two-tailed t-test; T(40) = -3.291, p = 0.002) without any alterations in the SW area (Fig. 7D; Mann-Whitney U test; p = 0.712), ripple amplitude (Fig. 7E; Mann-Whitney U test; p = 0.585) or ripple frequency (Fig. 7F; Mann-Whitney U test; p = 0.829).

**Fig 7.**
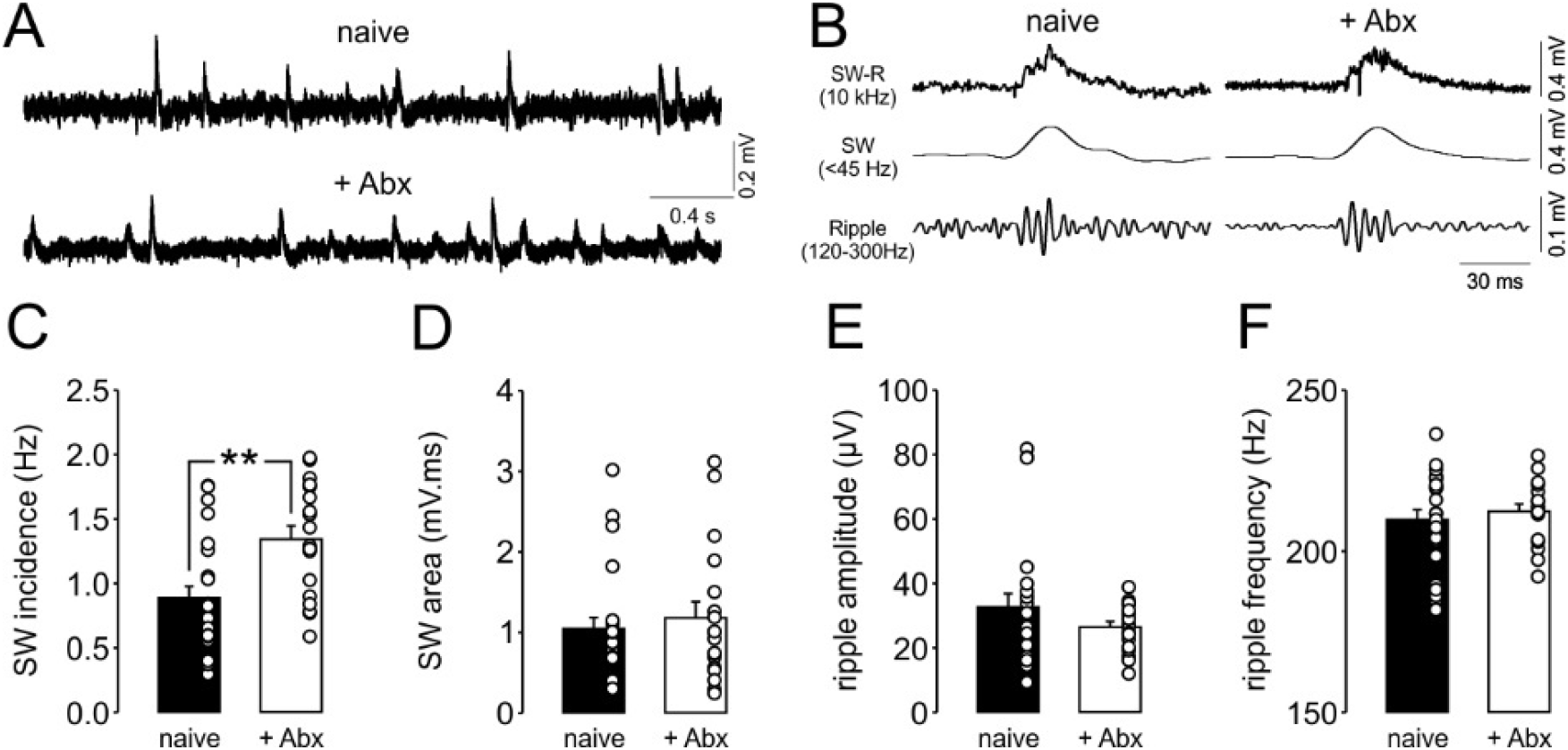
Increased incidence of sharp wave-ripples in the CA1 subregion of the hippocampus after chronic antibiotic treatment. **(A)** Representative LFP traces showing an increased number of sharp wave-ripples (SW-R) in the CA1 of +Abx (N=5 mice, n=18 slices for all parameters) in comparison to -Abx mice (N=7 mice, n=24 slices for all parameters). **(B)** Representative LFP traces of single SW-R, low-pass filtered SW (<45 Hz) and band-pass filtered ripples (120-300 Hz). Summary graphs showing **(C)** an increased CA1 SW incidence and no change in **(D)** SW area, **(E)** ripple amplitude and **(F)** ripple frequency upon chronic antibiotic treatment. Data are presented as mean ± SEM.

## 4. Discussion

In the current study, we demonstrate that alterations in homeostatic immunoregulation induced by gut dysbiosis is associated with impaired synaptic transmission and behaviorally-relevant brain rhythms in the hippocampus. We show that a four-week antibiotic treatment leads to a virtual depletion of the gut microbiota and substantial reduction in the circulating Ly6C^hi^ monocyte pool and decreased infiltration into the central nervous system (CNS). The reduced recruitment of peripheral cells was accompanied by increased activation of resident microglia with increased iNOS expression and reduced levels of the neurotrophin NGF and TNF. In line with the role of cytokines and neurotrophins in the maintenance of synaptic transmission, we observed a profound reduction of hippocampal CA3-to-CA1 synaptic transmission and cholinergic gamma oscillations. Thus, our study provides further insights into the interaction between gut microbiota and the CNS which is mediated by the immune system.

We have previously highlighted the role of circulating Ly6C^hi^ monocytes in gut-brain communication and their contribution to the recovery of Abx-induced impairment of hippocampal neurogenesis and cognitive function in mice (Möhle et al., 2016b). In the present study, we further investigated the consequences of the alterations in the circulating Ly6C^hi^ monocyte pool upon Abx-induced dysbiosis of the gut microbiota. We provide evidence for a shift towards an activated state of Ly6C^hi^ monocyte via enhanced expression of MHC II and CD40 (Fig. 2). A possible explanation for this observation could be an enhanced commensal translocation across intestinal epithelium and subsequent promotion of inflammation as we did observe enhanced proinflammatory gene expression of TNF and IL-6. This aligns with previous studies that described increased IL-17 and IFNγ after Abx-induced bacterial translocation or enhanced IL-6 and TNF in response to local bacterial infection in the spleen (Knoop et al., 2016; Straub et al., 2000).

Peripheral inflammation is associated with altered proinflammatory cytokine levels and circuit excitability in the brain (Galic et al., 2012). To gain insights into the potential regulation of proinflammatory cytokines in the CNS upon Abx treatment, we assessed the expression levels of TNF, IL-6 and IFNγ in whole brain homogenates (Fig. 3). Surprisingly, we found a moderate reduction in the TNF expression. In the brain, TNF is primarily synthesized in non-neuronal cell populations, including microglia and astrocytes (McCoy and Tansey, 2008). In contrast to the reduced TNF expression, we found a reduction in microglia-specific CX3CR1 expression, whereas we did not observe an altered GFAP expression (Fig. 4). Furthermore, we detected increases in markers that would suggest an activated microglial phenotype. Given the global reduction of TNF in the brain and the only basal infiltration of peripheral immune cells, it is likely that the observed increase in the percentage of TNF-producing microglia has only minimal influence on higher brain functions. While TNF has neuromodulatory effects by influencing neurotrophin production at low concentrations (Perry et al., 2002), high TNF concentrations are linked to neurotoxicity (Probert, 2015). Thus, prolonged Abx treatment could presumably increase TNF levels and result in detrimental effects. In line with the concept, a reduction in CX3CR1 expression or reduced interaction with its ligand CX3CL1 is associated with increased microglia activation and overproduction of pro-inflammatory cytokines (Cardona et al., 2006; Lyons et al., 2009; Rogers et al., 2011). Moreover, CX3CR1^-/-^ mice show deficits in hippocampal synaptic physiology and reduced coherence of distinct LFP network activities, including gamma oscillations (Rogers et al., 2011; Zhan et al., 2014). These observations are consistent with the involvement of activated microglia as mediators of reduced hippocampal synaptic transmission and gamma oscillatory activity after prolonged Abx treatment.

Reduced glial TNF production and subsequent signaling in neurons leads to impairment of excitatory synaptic transmission and synaptic scaling via regulation of surface AMPA-R levels (Beattie et al., 2002; Stellwagen and Malenka, 2006). In our experiments, Abx treatment lead to reduced expression of TNF and reduced hippocampal CA1 synaptic strength upon (Fig. 5), known to be dependent on intact AMPA-R function (Chater and Goda, 2014). However, transcriptional alterations of two main AMPAR subunits (GluR1, GluR2), the astrocytic (EAAT2), vesicular glutamate transporter (VGLUT1) and inhibitory GABA receptor (GABA_A_α1) which play an important role in sustaining synaptic transmission and plasticity in the brain, were not detected (Fig. 4). Thus, future studies assessing the cell-surface expression levels of AMPA-R in a brain-region specific manner might help in gaining more mechanistic insights into the impact of Abx treatment on circuit excitability.

Microglia activation can lead to increased generation of reactive oxygen species (ROS) and reactive nitrogen species (RNS) (Ramalingam and Kim, 2012). Their imbalanced production is associated with excessive oxidative stress which can be detrimental to cellular function and is commonly observed during many pathological conditions (Dröge and Schipper, 2007; Okuno et al., 2005; Sesti et al., 2010). Of note, cholinergic gamma oscillations appear to require high levels of mitochondrial activity and use high levels of oxidative capacity (Kann et al., 2011). These features make gamma oscillations especially vulnerable to oxidative stress (Hasam-Henderson et al., 2018). While gene expression levels of ROS producing enzymes NADPH-oxidase (*Nox4*) or Cyclooxygenase 2 (*Cox2*) were not altered, we measured a significant increase in the expression of iNOS (*Nos2*) (Fig. 4). This finding is particularly interesting in the light of the recent reports demonstrating that iNOS-mediated NO release can modulate cholinergic hippocampal gamma oscillations *in vitro* (Papageorgiou et al., 2016; Ta et al., 2019) and open a new perspective for synergistic interactions between microglia/microglia-associated factors and sustainment of gamma oscillations in the context of disease pathology (Adaikkan and Tsai, 2020; Iaccarino et al., 2016; Martorell et al., 2019). Future studies investigating specific microglial factors and their relation to intrinsic oscillatory brain activities in a brain-region specific manner will further help to understand the impact of long-term Abx treatment on distinct brain functions.

Gut dysbiosis can have major impacts on CNS function that results in altered cognitive function, mood and behavior (Irwin and Miller, 2007; Mayer, 2011; Sarkar et al., 2018). Converging evidence from our previous studies and by other groups indicate that Abx treatment leads to a distinct alteration memory, as evidenced by the novel object recognition test (Desbonnet et al., 2015; Fröhlich et al., 2016b; Möhle et al., 2016b; Sarkar et al., 2020). In addition to the previously described reduced hippocampal neurogenesis as a correlate of this memory deficit (Möhle et al., 2016b), now, we found a strong reduction in CBh-induced cholinergic gamma oscillations in the hippocampus upon Abx treatment (Fig. 6). In accordance, increase in CA3 gamma power and gamma coherence along the hippocampal CA3-CA1 axis appears to be critical for novel object recognition/exploration (Trimper et al., 2014). Furthermore, deficit in object place recognition task in an Alzheimer’s mouse model can be rescued by optogenetic gamma stimulation (Etter et al., 2019). Thus, an insufficient CA3 gamma power might underlie the deficit in this hippocampus-dependent memory. In support of this argument, we describe a mild but significant reduction in, which is known to positively regulate the cholinergic activity in the hippocampus (Conner et al., 2009). Furthermore, a recent study (Sarkar et al., 2020) provides evidence for a substantial elevation in brain ACh esterase levels upon Abx treatment. Such an increase in the ACh esterase might reduce extracellular ACh concentration and potentially diminish cholinergic gamma oscillations (Hollnagel et al., 2015). Interestingly, chronic Abx treatment leads to an aberrant sleep/awake architecture and a mild reduction in theta power during sleep suggesting a potential reduction of co-occurring gamma oscillations *in vivo* (Ogawa et al., 2020). Thus, we propose that, upon Abx treatment, a potential decline in the cholinergic tonus associated with reduced levels of NGF might contribute to the impairment of cholinergic gamma oscillations.

The duration of Abx-administration appears to be critical for determining the impact of gut dysbiosis on emotional behavior. While relatively short periods of Abx treatment (∼1 w) had no apparent effects on innate anxiety (Fröhlich et al., 2016b), longer Abx treatment (∼7 w) (Desbonnet et al., 2015) from weaning on reduced anxiety-like behavior in mice. Similarly, germ-free mice show also reduced anxiety-like behavior (Heijtz et al., 2011; Neufeld et al., 2011) resembling the behavioral alterations observed with longer periods of Abx treatment. Our electrophysiological recordings were obtained from slices obtained from the ventral-to-mid hippocampus. The ventral sector of the hippocampus has been implicated in mediating anxiety (Fanselow and Dong, 2010) and its lesion leads to reduced innate anxiety (Kjelstrup et al., 2002). Thus, a reduced baseline transmission and diminished gamma oscillations in the ventral hippocampus align well with a reduced anxiety and might be involved in the anxiolytic effects reported for long-term Abx treatment.

In addition to gamma oscillations, sharp wave-ripples (SW-R) are also generated in the CA3-to-CA1 axis of the hippocampus and the balance between these network activities is highly dependent on cholinergic tonus (Buzsáki, 2015; Çalışkan et al., 2016; Çalışkan and Stork, 2019). We found a profound increase in the incidence of SW-R upon long-term Abx treatment (Fig. 7), similar to the findings of a previous study demonstrating an increased hippocampal SW incidence in association with retarded extinction of contextual fear memories (Çalişkan et al., 2016). Accordingly, a recent work (Chu et al., 2019) shows that Abx-treated mice show impaired extinction of a cued fear memory. Both of these fear paradigms are dependent on the functional interactions between the hippocampus and amygdala (Çalışkan et al., 2019) and amygdalar activity appears to be recruited during SW events (Girardeau et al., 2017). Thus, increased SW incidence might partially predispose for re-consolidation of original fear memories and impair their extinction in Abx-treated mice.

To the best of our knowledge, this is the first study demonstrating the alterations in the LFP hippocampal network oscillations and synaptic properties of the hippocampus after long-term Abx treatment in mice. These electrophysiological alterations are associated with an enhanced activation of peripheral immune cells and microglia as well as alterations in the expression of neurotrophic factors and cytokines in the brain. A very recent study on mice raised under GF conditions from birth onwards, has demonstrated a reduced LTP in the hippocampal CA1 of male GF mice while no alteration was detected in the baseline synaptic transmission (Darch et al., 2021). As we found a profound reduction in the baseline synaptic transmission but only a tendency for a reduction in the CA1 LTP (Fig. 5), a functional microbiome during the early stages of life appears to be important for the sustainment of hippocampal plasticity during adulthood. Thus, studies targeting distinct developmental stages and adulthood are needed to elucidate the impact of Abx treatment on synaptic and cognitive function in an age-dependent manner. Further studies that assess more defined brain areas using causal intervention methods will increase our understanding of the brain-region specific impact of Abx treatment. The observed electrophysiological and gene expression changes after depletion of gut flora may have broader implications when considering psychiatric and neurodegenerative conditions associated with specific alterations in the immune homeostasis in humans.

## Acknowledgements

We are grateful to A. Koffi von Hoff, F. Blitz, G. Reifenberger, D. Zabler and S. Stork for excellent technical assistance and to A. Bohnstedt and D. Al-Chackmakchie for excellent animal care. We are grateful to Dr. J.O. Hollnagel for supporting us with the MATLAB-based analysis tools.

